# Hemocyte-targeted gene expression in the female malaria mosquito using the *hemolectin* promoter from *Drosophila*

**DOI:** 10.1101/2019.12.13.875518

**Authors:** Emilie Pondeville, Nicolas Puchot, Jean-Philippe Parvy, Guillaume Carrissimo, Mickael Poidevin, Robert M. Waterhouse, Eric Marois, Catherine Bourgouin

**Affiliations:** CNRS Unit of Evolutionary Genomics, Modeling, and Health (UMR2000), Institut Pasteur, Paris, France; Centre de Génétique Moléculaire, CNRS UPR 2167, Gif-sur-Yvette, France; Department of Ecology and Evolution, and Swiss Institute of Bioinformatics, University of Lausanne, 1015 Lausanne, Switzerland; CNRS UPR9022, INSERM U1257, Université de Strasbourg, Strasbourg, France

## Abstract

Hemocytes, the immune cells in mosquitoes, participate in immune defenses against pathogens including malaria parasites. Mosquito hemocytes can also be infected by arthropod-borne viruses but the pro- or anti-viral nature of this interaction is unknown. Although there has been progress on hemocyte characterization during pathogen infections in mosquitoes, the specific contribution of hemocytes to immune responses and the hemocyte-specific functions of immune genes and pathways remain unresolved due to the lack of genetic tools to manipulate gene expression in these cells specifically. Here, we used the Gal4-UAS system to characterize the activity of the *Drosophila* hemocyte-specific *hemolectin* promoter in the adults of *Anopheles gambiae*, the malaria mosquito. We established an *hml*-Gal4 driver line that we further crossed to a fluorescent UAS responder line, and examined the expression pattern in the adult progeny driven by the *hml* promoter. We show that the *hml* regulatory region drives hemocyte-specific transgene expression in a subset of hemocytes, and that transgene expression is triggered after a blood meal. The *hml* promoter drives transgene expression in differentiating prohemocytes as well as in differentiated granulocytes. Analysis of different immune markers in hemocytes in which the *hml* promoter drives transgene expression revealed that this regulatory region could be used to study phagocytosis as well as melanization. Finally, the *hml* promoter drives transgene expression in hemocytes in which o’nyong’nyong virus replicates. Altogether, the *hml* promoter constitutes a good tool to drive transgene expression in hemocyte only and to analyze the function of these cells and the genes they express during pathogen infection in *Anopheles gambiae*.

## Introduction

The mosquito *Anopheles gambiae* is the main African vector of *Plasmodium falciparum*, responsible for the severe forms of human malaria, which caused about 405 000 deaths in 2018 (WHO 2019). In the absence of an efficient vaccine, control of mosquito vectors remains the most important tool to fight malaria. Indeed, malaria vector control interventions based on insecticide-treated bed nets and indoor residual spraying were responsible for 78% of the decline in malaria cases between 2000 and 2015 (Bhatt, Weiss et al. 2015). However, these advances are hampered by the development of insecticide resistance by mosquitoes (Churcher, Lissenden et al. 2016, Ranson and Lissenden 2016). The failure of existing methods for eradicating malaria as well as advances in the field of vector genomics and genetics including mosquito genome editing with notably the recent advent of CRISPR-Cas9 gene drive technology in mosquitoes have sparked interest in genetic vector control strategies, including replacing wild populations with malaria refractory mosquitoes (McLean and Jacobs-Lorena 2016, Macias, Ohm et al. 2017). This latter approach consists of introducing genes into the mosquito genome so the mosquitoes become resistant to parasite infection thereby blocking malaria transmission. To this aim, the identification of promoters for directing transgene expression is crucial as transgene expression should be limited to a specific tissue and time in the mosquito to achieve maximum effect during pathogen exposure while minimizing the fitness cost on the vector (Terenius, Marinotti et al. 2008, Macias, Ohm et al. 2017). Importantly, effector genes conferring resistance to the parasite must also be identified. As mosquito immunity influences mosquito vector competence for malaria parasites (Clayton, Dong et al. 2014, Saraiva, Kang et al. 2016, Bartholomay and Michel 2018, Kumar, Srivastava et al. 2018), *i.e.* the ability to become infected following an infectious blood meal and to subsequently transmit the parasite, some of the proposed antimalarial strategies could be based on amplifying mosquito immune defenses.

Mosquito innate immunity is principally composed of two arms: i/ the humoral immune response including a complement-like system as well as the production of antimicrobial peptides (AMPs) and ii/ a cellular immune response mediated by hemocytes, the immune cells of insects (Clayton, Dong et al. 2014, Saraiva, Kang et al. 2016, Bartholomay and Michel 2018, Kumar, Srivastava et al. 2018). Mosquito hemocytes, which circulate in the hemolymph or are attached to tissues (sessile hemocytes), participate directly in immune defenses by phagocytizing pathogens in the hemolymph, or indirectly by secreting effector molecules mediating pathogen killing *via* lytic and melanization pathways (Blandin and Levashina 2007, Hillyer 2010, Hillyer and Strand 2014, Bartholomay and Michel 2018). In *An. gambiae* adults, three different types of hemocytes have been described based upon their morphology, granulocytes, oenocytoids and prohemocytes (Castillo, Robertson et al. 2006). Granulocytes, which are the most abundant cell type, are large cells with numerous granules in the cytoplasm. They are capable of phagocytosis and they can bind and spread on solid surfaces, forming filopodia and focal adhesions. Oenocytoids measure approximatively 9 µm in diameter with a homogeneous cytoplasm. They show phenoloxidase activity (involved in melanization) and are non-phagocytic. Prohemocytes are round and small cells of approximatively 4 to 6 µm with a high nuclear to cytoplasm ratio and are thought to be the putative hemocyte progenitor cells with the capacity to differentiate into other cell types. In mosquitoes, little is known about the origin of adult hemocytes. In *Drosophila*, adult hemocytes consist of embryonic hemocytes and larval hemocytes, which ones are released from the lymph glands at the onset of metamorphosis (Holz, Bossinger et al. 2003). While it was previously considered that adult flies do not have a hematopoietic organ, it is an ongoing matter of debate with a recent study showing that they possess active hematopoietic sites in the abdomen (Ghosh, Singh et al. 2015) and another study finding no increase in hemocyte numbers during adult life (Bosch, Makhijani et al. 2019). Although no hematopoietic organ has yet been identified in mosquitoes, hemocyte numbers increase after a blood meal (Castillo, Brown et al. 2011, Bryant and Michel 2014, Bryant and Michel 2016) and upon infection with bacteria or parasites (King and Hillyer 2012, King and Hillyer 2013, Ramirez, Garver et al. 2014). This increase in hemocyte number is thought to occur by mitosis of circulating hemocytes (King and Hillyer 2013, Bryant and Michel 2014).

In *An. gambiae* adult females, bacterial or parasite challenges also trigger hemocyte differentiation (King and Hillyer 2012, King and Hillyer 2013, Ramirez, Garver et al. 2014) and this is associated with differences in their gene and protein expression (Baton, Robertson et al. 2009, Pinto, Lombardo et al. 2009, Smith, King et al. 2016). Moreover, hemocytes are involved in phagocytosis (Moita, Wang-Sattler et al. 2005, King and Hillyer 2012, King and Hillyer 2013, Lombardo, Ghani et al. 2013) and in immune responses against malaria parasites (Pinto, Lombardo et al. 2009, Ramirez, Garver et al. 2014, Smith, Barillas-Mury et al. 2015, Lombardo and Christophides 2016, Smith, King et al. 2016). Immune cell differentiation has also been implicated in innate immune memory (Ramirez, Garver et al. 2014, Ramirez, de Almeida Oliveira et al. 2015, Barletta, Trisnadi et al. 2019). Hemocytes can be infected by o’nyong’nyong virus (ONNV), an arthropod-borne virus (arbovirus) transmitted by *An. gambiae* (Parikh, Oliver et al. 2009, Carissimo, Pondeville et al. 2015) but the pro- or anti-viral nature of this interaction is largely unknown. Although there has been progress on their characterization during pathogen infections in mosquitoes thanks to the rise of omics approaches and RNA interference-based gene silencing, the specific contribution of hemocytes to immune responses remains unclear. Recently, the role of granulocytes in anti-bacterial and anti-plasmodial immune responses has been elegantly demonstrated by chemical depletion of phagocytic cells (Kwon and Smith 2019). However, the hemocyte-specific functions of immune genes and pathways in mosquitoes are still unresolved due to the lack of genetic tools to manipulate gene expression in these cells specifically.

The Gal4-UAS system provides a perfect tool to characterize the expression pattern of promoters but also to study gene function at the tissue level. Indeed, this system (and its numerous extensions), established in the fly *Drosophila melanogaster* by Brand and Perrimon (Brand and Perrimon 1993) and routinely used since, has proven to be one of the most powerful techniques for addressing gene function *in vivo* (Duffy 2002, Southall, Elliott et al. 2008). This system relies on two components: Gal4, a transcriptional activator from yeast, which is expressed in a tissue-specific manner by the promoter placed upstream, and a transgene under the control of the upstream activating sequence (UAS) activated through Gal4 binding. The two components can be brought together using simple genetic crosses. This system is highly flexible, providing a versatile tool for controlling ectopic expression both spatially and temporally. Indeed, the expression of Gal4 can be controlled in a spatial and temporal manner using specific promoters, thereby dictating where the UAS-transgene is expressed. A key advantage of the system is the separation of the two components in different transgenic parental lines, which ensure that the transgene is silent until Gal4 is introduced in a genetic cross allowing transgenic lines encoding toxic or lethal proteins to be engineered. Moreover, a single UAS-transgene can be analyzed using different Gal4 drivers and different UAS-transgenes can be analyzed in the same tissue using a single Gal4 line, avoiding the creation of a new line for each promoter-gene combination. The Gal4-UAS system has since been developed in some model and pest species including *Arabidopsis* (Guyer, Tuttle et al. 1998), Zebrafish (Scheer and Campos-Ortega 1999), *Xenopus* (Hartley, Nutt et al. 2002), *Bombyx* (Imamura, Nakai et al. 2003) and *Tribolium* (Schinko, Weber et al. 2010). The Gal4-UAS system has been shown to be also functional in mosquitoes. Gal4 lines allowing specific expression in the midgut and in the fat body have been established in the arbovirus vector *Aedes aegypti* (Kokoza and Raikhel 2011, Zhao, Kokoza et al. 2014, Zhao, Hou et al. 2016). In *An. gambiae*, functional Gal4 lines have been established to drive transgene expression specifically in the midgut, in oenocytes or ubiquitously (Lynd and Lycett 2012, Adolfi, Pondeville et al. 2018, Lynd, Balabanidou et al. 2019). While hemocytes are important immune effectors, to date no hemocyte-specific Gal4 line has been established in mosquitoes. To develop such genetic tools, regulatory sequences need to be identified to drive gene expression in these cells only.

In *Drosophila*, hemocyte-specific transgene expression can be achieved using the *hemolectin* (*hml*) gene promoter (Goto, Kadowaki et al. 2003). Hml is a large protein similar to von Willebrand factor (Goto, Kumagai et al. 2001) which is produced by differentiated hemocytes only and is involved in coagulation and immunity against bacteria (Goto, Kadowaki et al. 2003, Lesch, Goto et al. 2007). Although different promoters can be used to drive gene expression in hemocytes in *Drosophila*, the *hml* promoter has the advantage that it drives expression only late during development (Goto, Kumagai et al. 2001, Goto, Kadowaki et al. 2003, Shia, Glittenberg et al. 2009) and genetic ablation of hemocytes driven by the *hml* promoter does not impede normal development (Shia, Glittenberg et al. 2009). Therefore, this promoter can be used to study gene function in mature hemocytes without the interference of indirect developmental effects. Here, we used the Gal4-UAS system to characterize the activity of the *Drosophila hml* promoter in *An. gambiae* adults. We established an *hml*-Gal4 driver line that we further crossed to a fluorescent UAS responder line we established previously (Adolfi, Pondeville et al. 2018), and examined the expression pattern in the adult progeny driven by the *hml* promoter. We show that the *hml* regulatory region drives hemocyte-specific transgene expression in a subset of hemocytes, and that transgene expression is triggered after a blood meal. *Hml* drives transgene expression in differentiating prohemocytes as well as in differentiated granulocytes. Analysis of different immune markers in hemocytes in which the *hml* promoter drives transgene expression revealed that this regulatory region could be used to study phagocytosis as well as melanization. Finally, the *hml* promoter drives transgene expression in hemocytes in which ONNV replicates. Therefore, this regulatory region could also be used to study the function of immune cells during an arboviral infection. Altogether, the *hml* promoter constitutes a good promoter to drive transgene expression in hemocyte only in order to analyze the function of these cells and the genes they express.

## Results

### Characterization of the expression pattern driven by the *Drosophila hml* promoter

The *hml*-Gal4 line was crossed with a fluorescent responder line UAS-mCD8::Cherry line we established previously (Adolfi, Pondeville et al. 2018), to characterize the expression pattern driven by the *hml* regulatory sequence in the adult progeny, referred as *hml*> mCD8::Cherry. To collect hemocytes, hemolymph was perfused from non-blood fed (NBF) females and blood fed (BF) females 24 hours post blood meal (PBM). Expression of *cherry* in hemocytes was analyzed by RT-PCR and by imaging. By RT-PCR, *gal4* was detected in hemocytes from both NBF and BF females, however *cherry* could only be detected in hemocytes from BF females (Figure 1, left panel). No *cherry* expression was detected in hemocytes collected from BF females from the control progeny of the crosses between the *hml*-Gal4 line or the UAS-mCD8::Cherry line to the E docking line (Figure 1, right panel), showing no leaking of the UAS construct.

**Figure 1.**
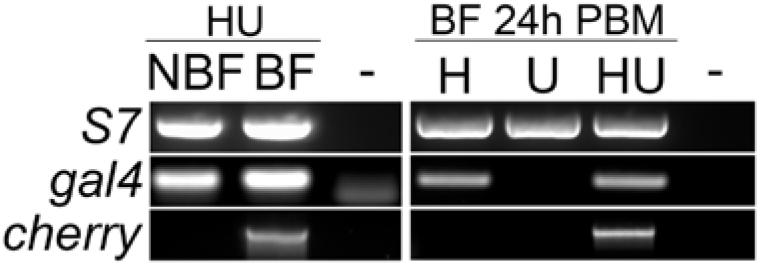
RT-PCR analysis of *gal4* and *cherry* expression in perfused hemocytes. Left panel: Expression of *gal4* and *cherry* in perfused hemocytes from non-blood fed (NBF) and blood fed (BF) females 24 hours post blood meal (PBM) issued from the cross between *hml*-Gal4 and UAS-mCD8::Cherry lines (HU females). Right panel: Expression of *gal4* and *cherry* in perfused hemocytes from BF females 24 hours PBM issued from the cross between *hml*-Gal4 and UAS-mCD8::Cherry lines (HU females) and from the control progeny of the crosses between the *hml*-Gal4 line or the UAS-mCD8::Cherry line to the E docking line, referred as H and U females. S7 is used as a standard gene. -: water negative PCR control.

Microscopy analyses confirmed the RT-PCR results. While we observed endogenous Cherry fluorescence in hemocytes collected from *hml*>mCD8::Cherry females 24h PBM (Figure 2), no Cherry fluorescence was detected in hemocytes perfused from the control progeny of the crosses between the *hml*-Gal4 line or the UAS-mCD8::Cherry line to the E docking line 24h PBM (data not shown). Perfused hemocytes can rapidly bind to the glass surface, and start to differentiate into granulocytes with filopodia. Therefore, perfused hemocytes from BF females were fixed either 10 or 45 min after the perfusions to analyze Cherry fluorescence at different hemocyte differentiation stages, *e.g.* prohemocytes and differentiated hemocytes. The hemolymph fixed 10 min after perfusion contained a large proportion of prohemocytes, (cell diameter of 5 to 6 µm) starting to differentiate and to produce filopodia (Figure 2A) as well as relatively small granulocytes with filopodia (Figure 2B-D). After 45 min on the slide, no prohemocyte could be observed and most of hemocytes were differentiated into large granulocytes, some of which measured up to 40 µm (Figure 2E-L). Probably due to their lack of adhesion (Castillo, Robertson et al. 2006), only few oenocytoids were observed and none of them displayed Cherry fluorescence. Cherry was present in both differentiating prohemocyte and granulocyte cell populations, although not in all cells from each population suggesting different cell lineages exist with similar morphology (Figure 2). Rarely, we also observed some giant rounded hemocytes expressing Cherry (Figure 2F and G), which could correspond to the “rounded” hemocytes described by King and Hillyer (King and Hillyer 2013). Interestingly, some large granulocytes contained two nuclei (*e.g.* Figure 2E and H), which are likely hemocytes in mitosis as previously reported (King and Hillyer 2013).

**Figure 2.**
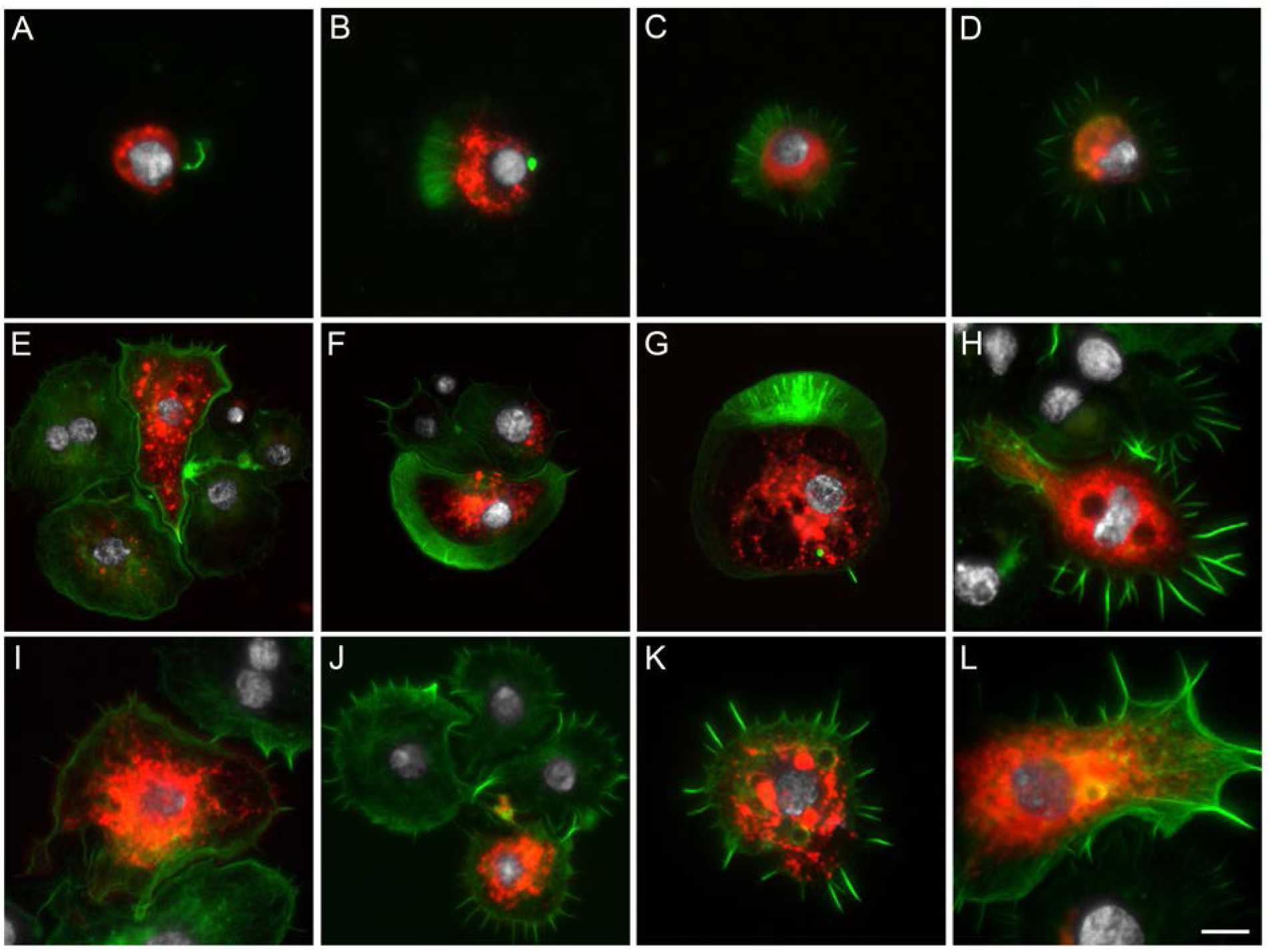
Microscopy analysis of hemocytes collected from *hml*>mCD8::Cherry blood fed females. Hemocytes expressing mCD8::Cherry under the control of *hml* promoter. (A-D) Hemocytes fixed 10 minutes post perfusion, (E-L) Hemocytes fixed 45 minutes post perfusion. Red: endogenous mCD8::Cherry; Green: Phalloidin; White: DAPI staining. Scale bar is 5 µm except for E, F and G where it is 10 µm.

In hemocytes expressing Cherry, the fluorescent protein was localized heterogeneously in the cytoplasm, most often in granules and vesicles. In the UAS responder line, Cherry is fused with mCD8, a transmembrane protein used as a cell membrane marker. Therefore, structures revealed by the membrane-bound Cherry are likely to be intracellular organelles such as endocytic vesicles. The cell surface staining around the filopodia of granulocytes was visible using high levels of exposure for picture acquisition (Figure S1A and B). Anti-mCD8 staining also confirmed that mCD8 and Cherry were exclusively expressed in the same hemocytes (Figure S1C and D). Cherry and mCD8 did not always colocalize within positive cells since we could observe Cherry but no mCD8 signal or mCD8 but no Cherry signal likely due to a cleavage of the fusion protein.

By imaging, very few hemocytes expressing Cherry were seen in perfusions either from NBF *hml*>mCD8::Cherry females, confirming the RT-PCR results (Figure 1), or from *hml*>mCD8::Cherry males (data not shown) compared to BF *hml*>mCD8::Cherry females. In BF *hml*>mCD8::Cherry females, expression of the transgene under the *hml* promoter was activated after the blood meal, with a peak of positive hemocytes around 24-30h PBM and a decline to basal levels around 48-72h PBM. The proportion of perfused hemocytes which were positive varied greatly between experiments ranging roughly from 5 to 20% of total hemocytes from BF females. In rare cases and independently of the experimental context (pathogen or not), up to 60% of perfused hemocytes expressed the transgene. Some hemocytes also expressed low levels of Cherry and were only visible by microscopy when using high exposure levels. It is therefore possible that Cherry-positive hemocytes have been underestimated in some cases when visually observing the slides with the microscope objectives (*e.g.* Figure S2). As groups of 10 females were perfused per well, we wondered if the mix of Cherry-expressing and non-expressing hemocytes could be due to individual variation of transgene expression in hemocytes under the *hml* regulatory region. While there was variation in the proportion of Cherry-expressing hemocytes between perfusions from different individuals, every individual BF female contained both Cherry positive and negative hemocyte populations (data not shown).

In BF *hml*>mCD8::Cherry females at 24h PBM, Cherry-expressing hemocytes were also found attached to tissues, mostly close to the heart, abdominal muscles and epidermis (Figure 3A-C). These sessile hemocytes were differentiated with long filopodia which can be visualized with the mCD8::Cherry (Figure 3D). A few Cherry-positive hemocytes were also found attached to the midgut and the Malpighian tubules (Figure 3E and F). Cherry-expressing hemocytes were also found in close contact with tracheae (Figure 3G). As observed with hemolymph samples, the number of hemocytes expressing Cherry was variable between individuals with some showing a large number of positive hemocytes in some cases (Figure 3C). Imaging analysis of the tissues from *hml*>mCD8::Cherry females did not reveal other cells/tissues other than hemocytes expressing *hml*-driven Cherry fluorescence. Consistent with the perfusion results, very few hemocytes expressing Cherry were seen in the tissues from either NBF *hml*>mCD8::Cherry females or *hml*>mCD8::Cherry males compared to BF *hml*>mCD8::Cherry females and none in *hml*-Gal4 and UAS-mCD8::Cherry control females 24h PBM (data not shown).

**Figure 3.**
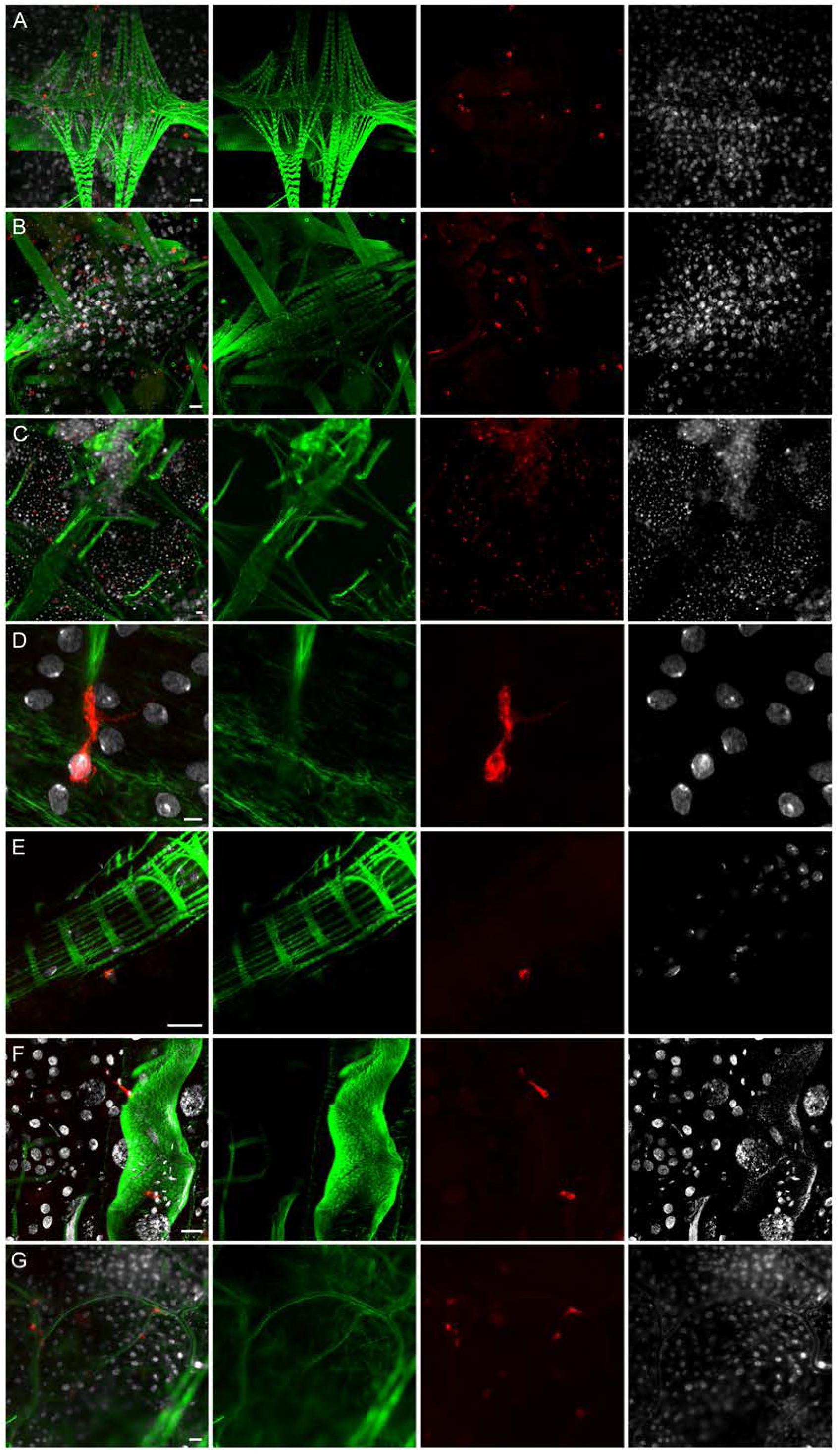
Microscopy analysis of tissues from *hml*>mCD8::Cherry blood fed females. Hemocytes expressing the mCD8::Cherry under the control of the *hml* promoter. (A,B,C) Abdominal muscles and epidermis, (D) Enlargement of a hemocyte attached to a muscle fiber, (E) Hemocyte attached to the anterior midgut, (F) Hemocytes attached to Malpighian tubule, (G) Hemocytes in close contact to tracheae. Red: endogenous mCD8::Cherry; Green: Phalloidin; White: DAPI staining. Scale bar is 20 µm except for D where it is 5 µm.

### Characterization of *hml*-driven Cherry-expressing hemocytes

To better characterize the hemocytes in which the *hml* promoter drives transgene expression, we then analyzed different immune markers in hemocytes from *hml*>mCD8::Cherry females. Granulocytes can phagocytose bacteria but also microspheres (Hillyer, Schmidt et al. 2003, King and Hillyer 2012). Therefore, hemocytes were perfused from *hml*>mCD8::Cherry females previously injected either with fluorescent microspheres (Figure 4A and B) or with a GFP-expressing *Escherichia coli* strain. Microscopy analysis showed that granulocytes expressing Cherry, as well as granulocytes which do not, were able to phagocytose bacteria and microspheres (Figure 4C and D). We also stained hemocytes perfused from BF *hml*>mCD8::Cherry females with an antibody against the thioester containing protein 1 (TEP1). TEP1 is a homolog of the mammalian C3 complement factor secreted in the hemolymph where it can bind to malaria parasites, leading to their elimination through lysis and melanization (Levashina, Moita et al. 2001, Blandin, Shiao et al. 2004). In case of a bacterial challenge, TEP1 protein binds to bacteria and is taken up by hemocytes, most likely through phagocytosis of TEP1-tagged bacteria (Levashina, Moita et al. 2001, Moita, Wang-Sattler et al. 2005, Volohonsky, Hopp et al. 2017). We detected TEP1 in the cytoplasm of some granulocytes, expressing Cherry or not (Figure 5). In granulocytes in which the *hml* promoter drives expression of the fluorescent protein, the mCD8 membrane marker revealed that TEP1 was localized in vesicles very similar to phagosomes described in *Drosophila* hemocytes (Shandala, Lim et al. 2013). Phagosomes are intracellular vesicles formed after internalization of extracellular material by invagination of the plasma membrane and they interact with endosomes and lysosomes to form acidic phagolysosomes, which degrade pathogens. These phagosome-like structures were also observed in other perfusion experiments (*e.g.* Figure 6). Although we did not expose the females to a specific pathogen, mosquitoes were not reared in sterile conditions and the TEP1 staining in hemocytes could be due to phagocytosis of bacteria.

**Figure 4.**
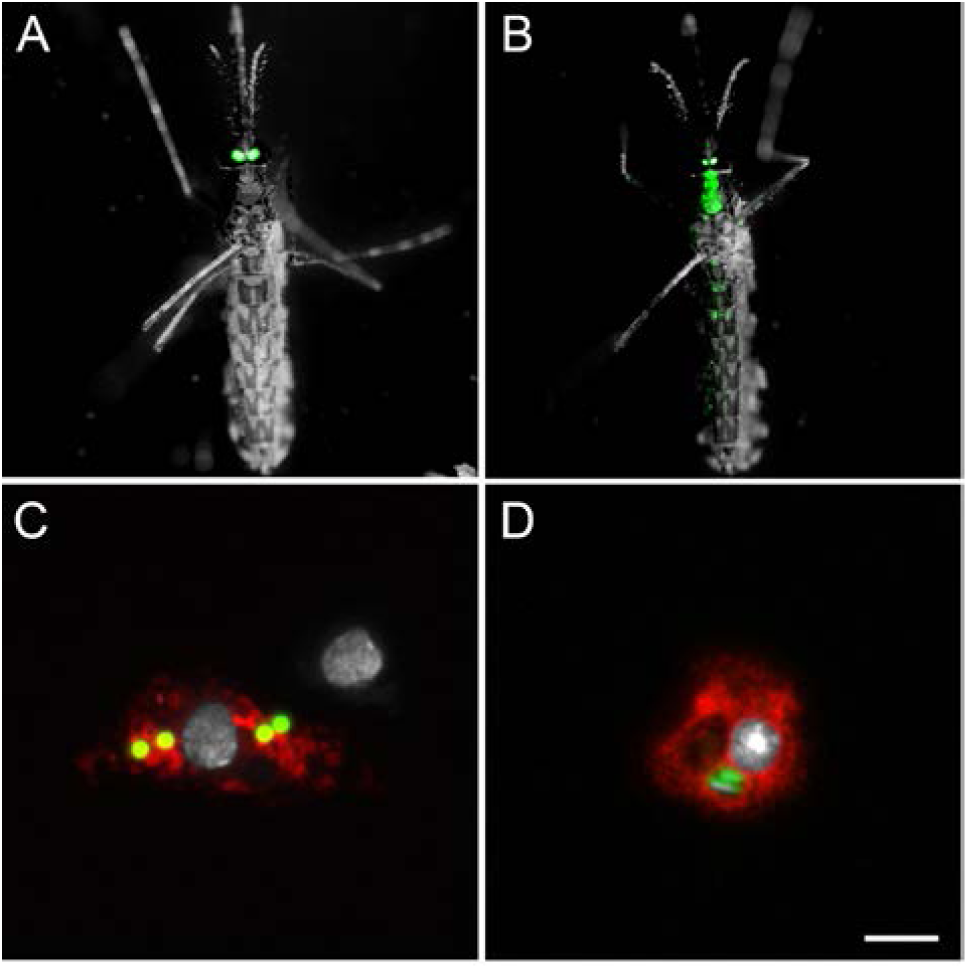
Hemocytes collected from *hml*>mCD8::Cherry females injected with fluorescent beads or bacteria. (A) *hml*>mCD8::Cherry control female showing eYFP expression in the eyes under the 3P3 promoter, (B) *hml*>mCD8::Cherry female injected with fluorescent beads, (C) Hemocytes collected from *hml*>mCD8::Cherry females injected with fluorescent beads, (D) Hemocytes collected from *hml*>mCD8::Cherry females injected with GFP-tagged *Escherichia coli*. Red: endogenous mCD8::Cherry; Green: fluospheres in B and C / *Escherichia coli* bacteria in D; White: DAPI staining. Scale bar is 5 µm.

**Figure 5.**
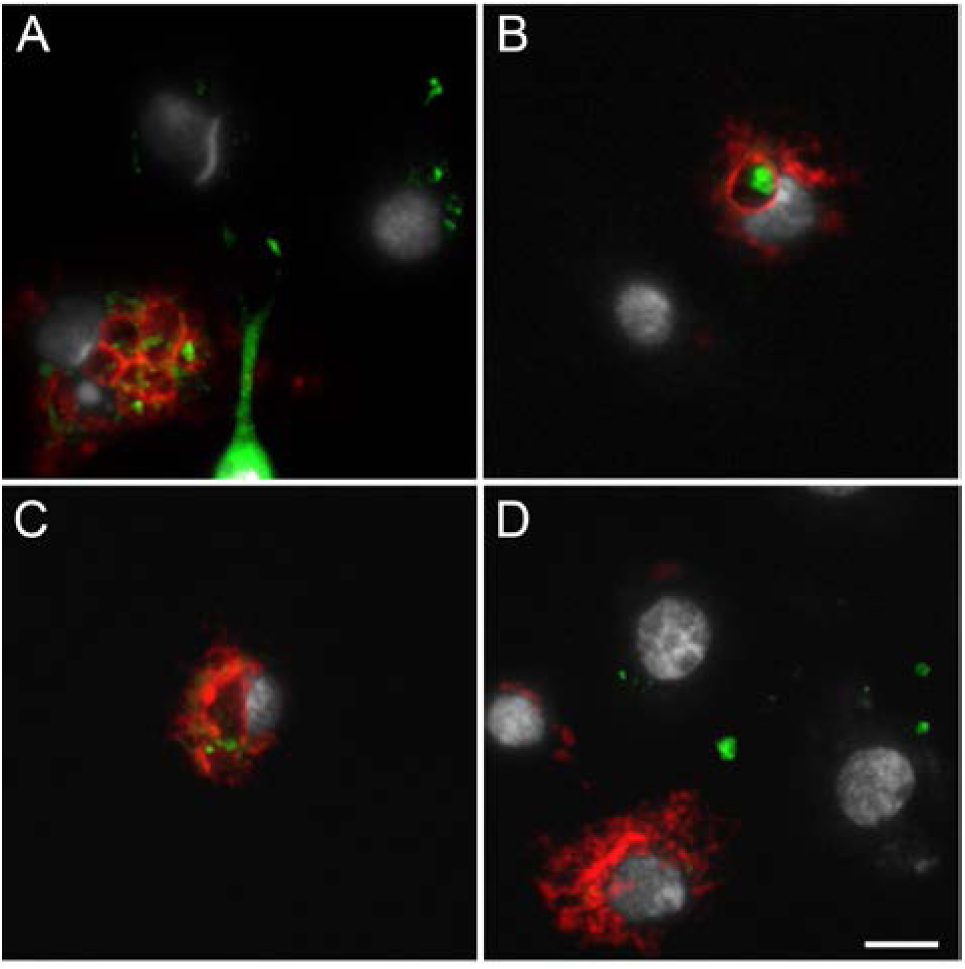
Anti-TEP1 immunostaining of hemocytes collected from *hml*>mCD8::Cherry blood fed females. Both the mCD8::Cherry under the control of *hml* promoter and TEP1 are detected in hemocytes (A, B, C), or only mCD8::Cherry (D), or only TEP1 (A, D), or none of the proteins (B). Red: endogenous mCD8::Cherry; Green: anti-TEP1 staining; White: DAPI staining. Scale bar is 5 µm.

**Figure 6.**
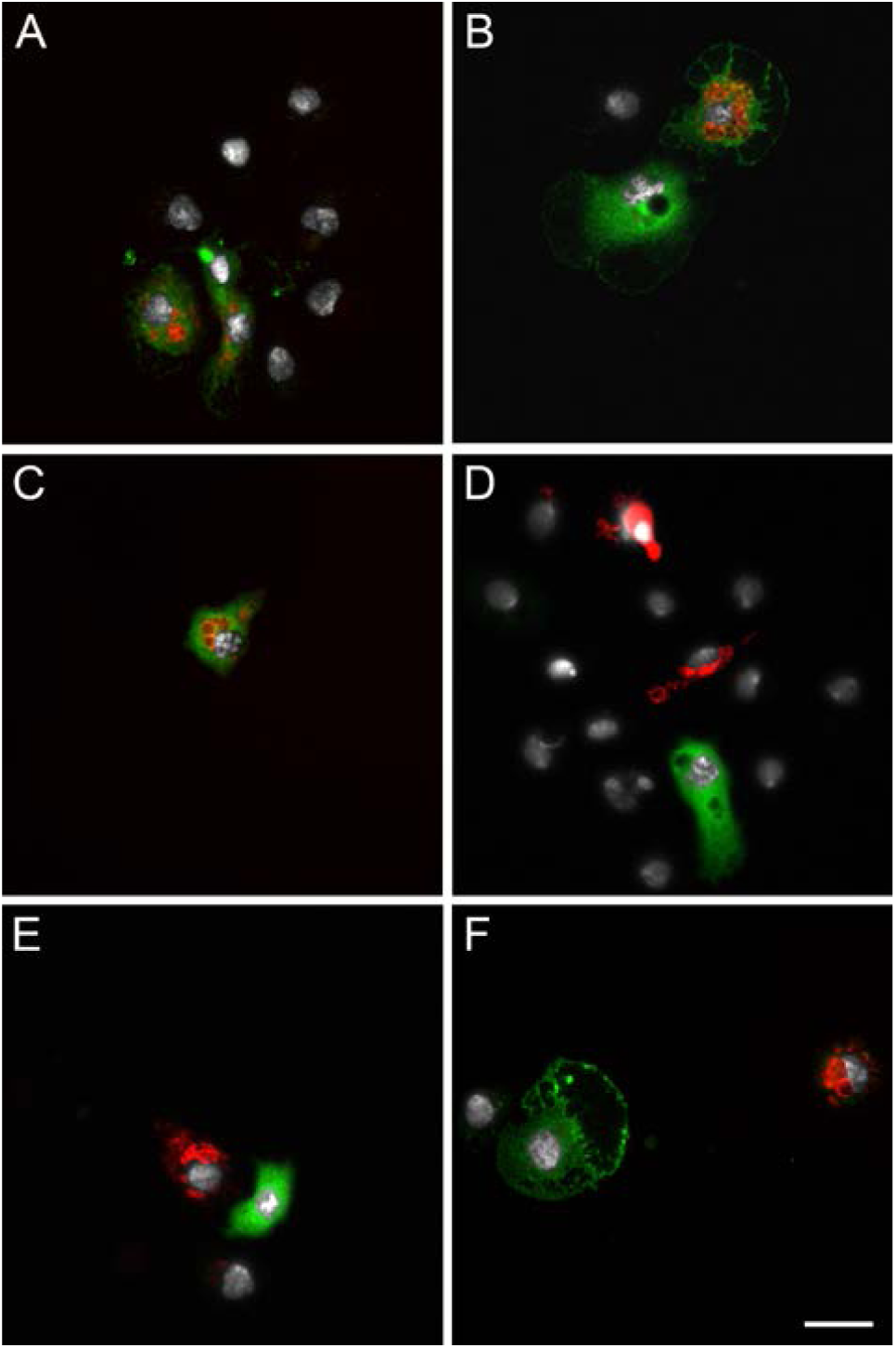
Anti-PPO2 immunostaining of hemocytes collected from *hml*>mCD8::Cherry blood fed females. Hemocytes express either both the mCD8::Cherry under the control of *hml* promoter and PPO2 (A, B, C), or only mCD8::Cherry (D, E, F), or only PPO2 (B, D, E, F), or none of the proteins (A,B,D,E,F). Red: endogenous mCD8::Cherry; Green: anti-PPO2 staining; White: DAPI staining. Scale bar is 10 µm.

In addition to phagocytosis, hemocytes also participate in melanization, a humoral immune response leading to the sequestration of invading pathogens into a thick proteinaceous layer. Hemocytes express phenoloxidases (POs), which are essential enzymes in the melanization cascade (Kumar, Srivastava et al. 2018). To determine if hemocytes in which the *hml* regulatory sequence drives transgene expression could be involved in the melanization process, we analyzed the expression of the prophenoloxidase 2 (PPO2) by immunostaining in perfused hemocytes from BF *hml*>mCD8::Cherry females. PPO2 is one of the ten *An. gambiae* PPOs and it is expressed in a hemocyte-like cell line (Muller, Dimopoulos et al. 1999) and in the hemolymph (Fraiture, Baxter et al. 2009). PPO2 was found in granulocytes and only in a subset of Cherry-expressing granulocytes. Three different granulocyte populations were observed, some hemocytes expressing both PPO2 and Cherry (Figure 6A-C), some expressing PPO2 or Cherry (Figure 6D-F) and some hemocytes expressing neither PPO2 nor Cherry. We also perfused hemocytes from adult transgenic mosquitoes expressing the tdTomato fluorescent protein under the control of the *An. gambiae* promoter of the *PPO6* gene (Volohonsky, Terenzi et al. 2015). As for PPO2 and the *hml*-driven transgene expression, PPO6-driven expression of tdTomato was found in only some granulocytes of BF females (Figure 7). While the *hml* promoter drives transgene expression in few hemocytes in NBF females and in males compared to BF females, the expression pattern driven by *PPO6* promoter in perfused hemocytes was similar between NBF females, BF females and males (NBF females and males not shown).

**Figure 7.**
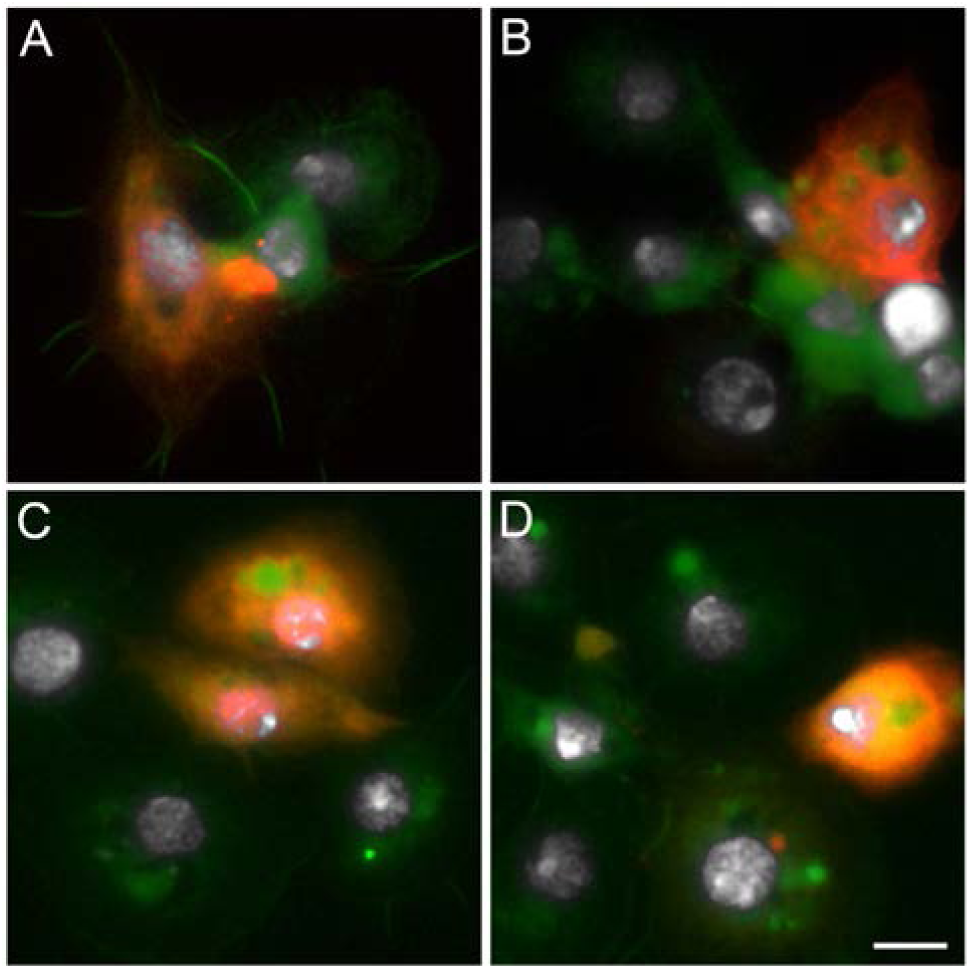
Hemocytes collected from *PPO6*-tdTomato mosquitoes. Red: endogenous tdTomato; Green: phalloidin; White: DAPI staining. Scale bar is 5 µm.

### Characterization of *hml*-driven Cherry-expressing hemocytes upon arbovirus infection

While most arthropod-borne viruses (arboviruses) transmitted by mosquitoes are transmitted by *Aedes* and *Culex* but not *Anopheles* mosquitoes, *An. gambiae* can be infected by, and transmit, ONNV (Williams, Woodall et al. 1965, Vanlandingham, Hong et al. 2005). Although the roles of hemocytes in mosquito antiviral immunity have not been extensively studied due to the lack of genetic tools, it has been shown that arboviruses can infect hemocytes in *Aedes aegypti* (Sindbis virus, (Parikh, Oliver et al. 2009)) and *An. gambiae* (ONNV, (Carissimo, Pondeville et al. 2015)). To assess if the *hml* promoter could drive transgene expression in hemocytes infected with ONNV, *hml*>mCD8::Cherry females were given either a non-infected blood meal or a blood meal infected with ONNV-eGFP (Brault, Foy et al. 2004). We perfused their hemocytes seven days later, a time at which ONNV has disseminated from the gut to other tissues and hemocytes (Carissimo, Pondeville et al. 2015). The GFP infection marker was expressed in hemocytes from infected females and some of the infected hemocytes expressed Cherry (Figure 8), showing that the *hml*-Gal4 line constitutes a valuable tool for investigating further the contribution of hemocytes to arbovirus infection in mosquitoes.

**Figure 8.**
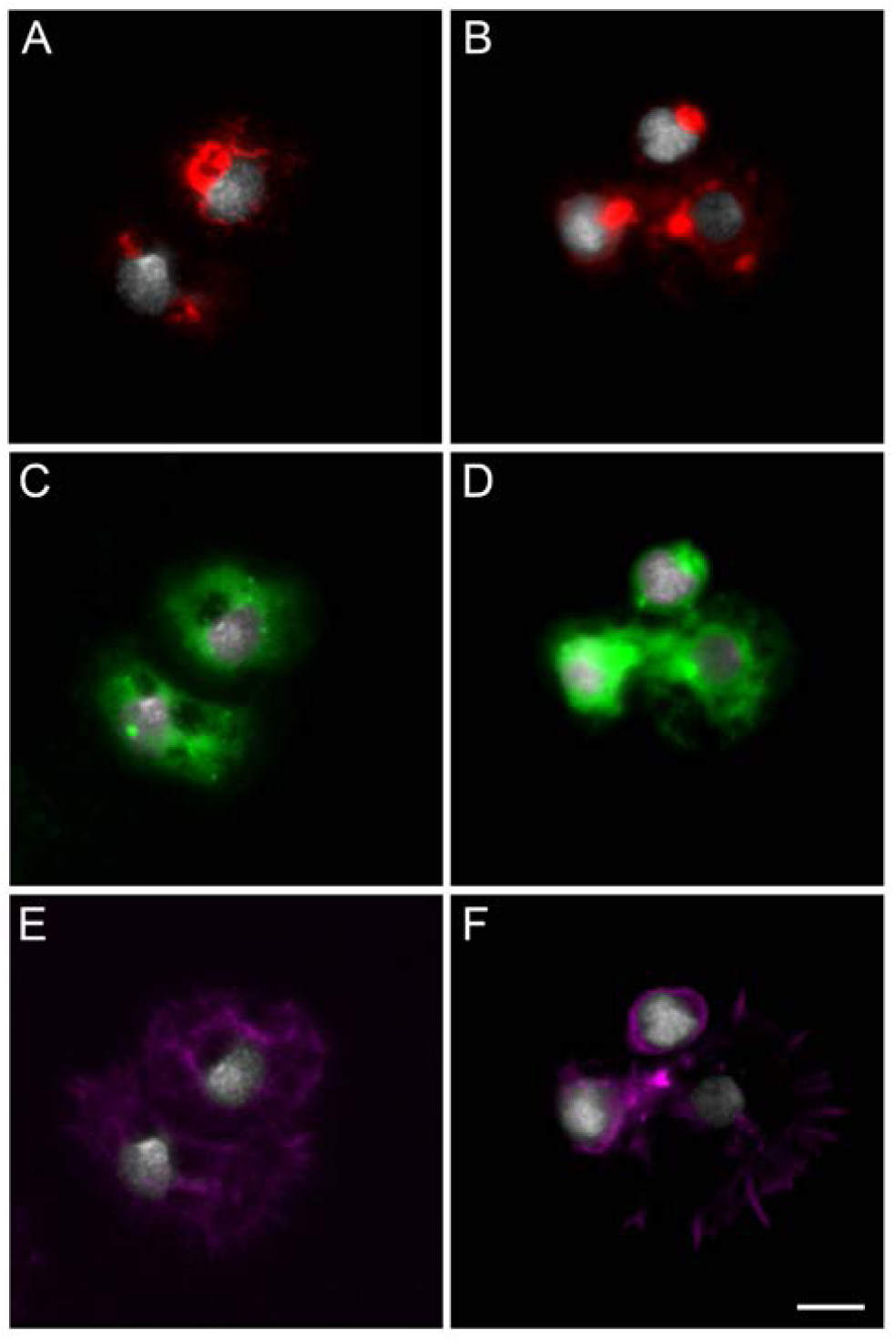
Hemocytes collected from *hml*>mCD8::Cherry females infected with o’nyong’nyong virus (ONNV). Red: endogenous mCD8::Cherry; Green: ONNV-eGFP; Magenta: phalloidin; White: DAPI staining. Scale bar is 5 µm.

## Discussion

Insect hemocytes play a fundamental role in insect immunity (Strand 2008, Hillyer and Strand 2014). However, their specific contributions in mosquitoes remain poorly understood due to the lack of genetic tools to study gene function in these specific cells. Here, we used the Gal4-UAS system to characterize the expression pattern driven by the *Drosophila hml* promoter in *An. gambiae* adult females. The *Drosophila hml* gene is homologous to the von Willebrand factor in vertebrates, and it is well conserved across animals including insects from different orders such as the lepidopteran *Bombyx mori* (Kotani, Yamakawa et al. 1995, Arai, Ohta et al. 2013) and in the hymenopteran *Apis mellifera* (Gabor, Cinege et al. 2017). At the time of this study design, we could not identify an orthologue of *hml* in the genome of *An. gambiae*, suggesting a putative gene loss event or the failure to sequence and assemble the locus encoding the *hml* orthologue. Subsequent progress in genome sequencing efforts, including from the *Anopheles* 16 genomes project (Neafsey, Waterhouse et al. 2015), has greatly improved genomic sampling of mosquito and other dipteran species. Intriguingly, no *hml* orthologues could be identified from any of the mosquito genomic resources at VectorBase (Giraldo-Calderon, Emrich et al. 2015), while clear orthologues are present in tsetse flies and the housefly (Figure S3), thereby strongly supporting the loss of the *hml* gene from the ancestral mosquito genome. Despite this relatively ancient loss of *hml*, our study reveals that the *Drosophila hml* promoter drives transgene expression in *An. gambiae*, showing that some mosquito transcription factors can still bind to this regulatory region and activate transgene expression.

Although the *hml* promoter could drive gene expression in a few hemocytes in males and NBF females, we found that transgene expression was mostly activated by blood feeding. We observed that transgene expression under the *hml* promoter peaked around 24h PBM and returned to pre-blood meal levels 48 hours after a blood meal. In *An. gambiae*, hemocytes respond to blood feeding by increasing their proliferation, size and granularity (Castillo, Robertson et al. 2006, Bryant and Michel 2014). Blood feeding also triggers a shift in proteins detected in hemocytes, including the immune factors TEP1, a complement-like molecule, and PPO6, which is involved in the melanization pathway (Bryant and Michel 2014, Smith, King et al. 2016). Similarly to the *hml*-driven gene expression, this hemocyte immune activation following a blood meal is transient with a return to pre-blood meal state within 48 hours (Bryant and Michel 2016). This suggests that blood feeding itself could prime the female immune system for potential blood-borne pathogen challenge. As Hml is involved in immunity in *Drosophila* (Lesch, Goto et al. 2007) and is a marker of differentiated hemocytes (Goto, Kadowaki et al. 2003, Evans, Liu et al. 2014), it is therefore not surprising that the *hml* promoter becomes active after blood feeding in a similar way to other mosquito immune genes expressed in granulocytes.

While hemocyte proliferation after a blood meal is controlled at least by the insulin signaling pathway (Castillo, Brown et al. 2011), factors mediating hemocyte activation after a blood meal are still unknown. This transient activation occurs simultaneously to an increase in Ras-MAPK signaling, suggesting that this pathway could be involved in hemocyte proliferation and differentiation (Bryant and Michel 2014, Bryant and Michel 2016). This could also be triggered by 20-hydroxyecdysone (20E) as the titers of this vitellogenic hormone, produced by ovaries after a blood meal, follow the same temporal pattern in mosquitoes including *An. gambiae* (Clements 1992, Pondeville, Maria et al. 2008). This latter hypothesis is further supported by the fact that blood feeding as well as 20E injection activates the expression of the Leucine-Rich repeat IMmune protein LRIM9, an antagonist of *Plasmodium berghei* development (Upton, Povelones et al. 2015). The expression of PPO1, another prophenoloxydase involved in melanization, is also up-regulated by 20E in an *An. gambiae* cell line (Ahmed, Martin et al. 1999). In *D. melanogaster*, steroid hormone signaling during metamorphosis in hemocytes is required for hemocyte immune functions and survival to bacterial infections (Regan, Brandao et al. 2013). Altogether, these findings suggest that mosquito female evolution towards anautogeny (requirement of a blood meal to develop eggs) co-opted molecules with functions in reproduction and whose expression is triggered by the blood meal to anticipate the risk of exposure to blood-borne pathogens. Others have hypothesized that this immune activation following a blood meal could be due to the leak of gut-resident bacteria into the hemolymph caused by the midgut epithelium stretch occurring during blood feeding (Bartholomay and Michel 2018). However, the peritrophic matrix, which is induced by the gut microbiota after a blood meal, normally prevents systemic dissemination of bacteria from the gut (Rodgers, Gendrin et al. 2017). Therefore, bacterial leak from the gut might be a rare event happening accidentally. Even though *hml*-driven expression in hemocytes was always induced in blood fed females, the proportion of positive hemocytes was much higher in rare cases for unidentified reasons. Therefore, it is possible that this rare and strong induction could be due to bacteria leaking from the gut. This would be consistent with Hml being involved in clotting and responses against bacteria as well as *hml* expression being induced by injuries in adult flies (Goto, Kumagai et al. 2001, Lesch, Goto et al. 2007).

In *Drosophila, hml* is only expressed in differentiating and differentiated hemocytes, and more precisely in crystal cells and plasmatocytes (Goto, Kadowaki et al. 2003, Lesch, Goto et al. 2007). Although they are named differently, the classification of hemocytes based on morphology, immune markers and function shows that crystal cells and plasmatocytes in flies correspond to oenocytoids and granulocytes respectively in mosquitoes (Castillo, Robertson et al. 2006). We also found that the *hml* promoter drives transgene expression in granulocytes. We did not detect oenocytoids in our experiments, probably because of their lack of adhesion. Therefore, we cannot exclude that the *hml* promoter could drive gene expression in these cell type as well. Although they had the same morphology, the *hml* promoter drove expression in a subset of granulocytes only, either in collected hemolymph or attached to tissues. As previously described, sessile hemocytes expressing Cherry were found mostly attached to the epidermis and abdominal muscles, in close contact to the tracheae and in periostal regions (King and Hillyer 2013). It was difficult to estimate an exact proportion of granulocytes expressing the transgene under the control of the *hml* promoter, due to variation between individuals and experiments, over the time after a blood meal, not only in the number but also in the intensity of the responder fluorescent protein. We did not identify the reason of this variation. However, as already observed and discussed for the *PPO6*-tdTomato line (Volohonsky, Terenzi et al. 2015), variation could be due to variegation effects. As mosquitoes were not reared in sterile conditions, it is also possible that some microbes influence *hml*-driven expression.

Our data showing that gene expression driven by the *hml* promoter in a subset of granulocytes, either in circulation or attached to tissues, is consistent with what has been described in *Drosophila* (Goto, Kadowaki et al. 2003, Shia, Glittenberg et al. 2009) and in *Apis mellifera* (Gabor, Cinege et al. 2017). Similarly, PPO2 was not expressed in all granulocytes, sometimes co-expressed with the transgene and sometimes not, highlighting at least four populations of granulocytes. Similar results were obtained with the *PPO6*-tdTomato line (Volohonsky, Terenzi et al. 2015, Severo, Landry et al. 2018) in which we detected transgene expression in some granulocytes only. Our data are in agreement with previous studies in *Anopheles* mosquitoes. Indeed, different immune markers such as Sp22D, PPO4 and PPO6 were expressed only in a subset of granulocytes (Castillo, Robertson et al. 2006, Bryant and Michel 2014, Severo, Landry et al. 2018, Kwon and Smith 2019). Therefore, different populations of granulocytes are likely to be present in *Anopheles* mosquitoes, even though they display the same morphology. Therefore, genetic analysis of immune cell functions as well as hemocyte-specific gene functions in mosquitoes might require the use of different regulatory regions to target the different subpopulations of hemocytes, as in *Drosophila* (Evans, Liu et al. 2014). Such regulatory regions could be identified by enhancer trapping, previously used in *An. stephensi* to identify salivary gland, midgut and fat body regulatory regions (O’Brochta, Pilitt et al. 2012). Transcriptomic and proteomic studies comparing gene and protein expression between hemocytes and carcass in *An. gambiae* identified some genes enriched in immune cells (Baton, Robertson et al. 2009, Lombardo, Ghani et al. 2013, Smith, King et al. 2016, Kwon and Smith 2019). Promoters of these genes could therefore constitute other good candidates to drive expression in mosquito hemocytes. However, this does not exclude that these genes are expressed at low levels in other tissues, or that they can be expressed in other tissues under different experimental conditions or during mosquito development (*e.g.* embryonic and larval development). For instance, transgene expression in *Drosophila* hemocytes can be achieved using the *hml* promoter, but also *peroxidasin* or *serpent* promoters. These two other promoters drive expression from embryonic development, resulting in lethality during development when combined with some transgenes, and can sometimes be non-specific to hemocytes driving the expression in the fat body, as observed for *peroxidasin* (Shia, Glittenberg et al. 2009).

One extension of the Gal4-UAS system to overcome this limitation is the use of the Temporal And Regional Gene Expression Targeting (TARGET) system developed in *Drosophila* (McGuire, Mao et al. 2004) and further established in the Zebrafish (Faucherre and Lopez-Schier 2011). In this system, flexible temporal and/or tissue control of Gal4 activity can be achieved by using the temperature-sensitive version of Gal80 protein (Gal80ts) under the control of tissue- or time-specific promoters. At permissive temperatures (18°C to 20°C), Gal80ts binds to Gal4 and inhibits Gal4 activity, while at restrictive temperatures (28 to 30°C), it no longer binds to Gal4, allowing Gal4-dependent transgene expression. This system would also be suitable for studying *Plasmodium falciparum* development in transgenic mosquitoes as the human malaria parasite, contrary to the rodent parasite, *Plasmodium berghei*, can develop within temperatures ranging from 21 up to 34°C, with most studies analyzing malaria infection in *Anopheles* at 24 to 28°C (Shapiro, Whitehead et al. 2017). The *hml*-gal4 line presented here was initially designed to carry, in tandem with *hml*-gal4, the gal80ts sequence under the control of a second and identical *hml* promoter. Although we could detect by RT-PCR the expression of *gal80ts* in perfused hemocytes, we could not observe any obvious difference in Cherry expression between blood fed mosquitoes reared at 20 versus 30°C (data not shown). Different hypotheses could explain this negative result. First, contrary to *D. melanogaster, An. gambiae* mosquitoes live and can cope with high temperatures. Therefore, the thermo-sensitivity of Gal80ts could be decreased in mosquitoes if chaperone proteins stabilize Gal80ts. Secondly, as the *hml* promoter does not drive expression in all hemocytes, with variation between individuals and experiments, and sometimes in a low proportion of granulocytes, it might be difficult to observe a significant decrease. Further studies will be required to assess the functionality of the Gal80ts system in mosquitoes. This could be undertaken by using an ubiquitous promoter such as the one of the *An. gambiae Polyubiquitin-c* (PUBc) gene (Adolfi, Pondeville et al. 2018).

In summary, the *hml* promoter combined with the Gal4-UAS system is a promising tool to study gene function in hemocytes, more particularly in a subpopulation of granulocytes, and their specific contributions to immunity in the malaria mosquito. Our findings show that it could be used to study blood meal-induced activation of hemocytes, phagocytosis, melanization as well as arbovirus-hemocyte interactions. Moreover, genes enriched in hemocytes and whose functions are yet unknown could be analyzed using this system. As phagocytosis of *Plasmodium* occasionally occurs (Hillyer, Schmidt et al. 2003) but is not believed to be required for an efficient anti-parasitic response in mosquitoes, we did not determine if granulocytes expressing the transgene under the *hml* promoter control could also phagocytose sporozoites after *Plasmodium* infection. Nevertheless, granulocytes are definitely involved in the response against the parasite (King and Hillyer 2012, Bartholomay and Michel 2018, Kwon and Smith 2019), suggesting that the *hml* promoter could also constitute a good driver for the analysis of hemocyte-mediated anti-malaria immunity. Some other fundamental questions, already raised by Hillyer and Strand (Hillyer and Strand 2014), about hemocyte origin lineage, function of hemocytes during post-embryonic development as well as the function of sessile hemocytes *in vivo*, could also be addressed using such genetic tools. The *gal4* sequence used in this study encodes the classic Gal4 form. Future studies could also assess whether transgene expression driven by the *hml* regulatory sequence can be increased by using Gal4 modified forms, for example Gal4Δ (Ma and Ptashne 1987) or Gal4GFY, shown to be more active than the classic Gal4 in an *An. gambiae* cell line (Lynd and Lycett 2011). Finally, it would be interesting to assess if the *hml* regulatory region from *Drosophila* could also be active in other mosquito species, including *Ae. aegypti*, the main mosquito vector of arboviruses.

## Material and methods

### Rearing of mosquito transgenic strains

The E phiC31 docking line (Meredith, Basu et al. 2011), the *hml*-gal4 (H) line, the UAS-mCD8::Cherry (U) line (Adolfi, Pondeville et al. 2018) and the *PPO6*-tdTomato line (Volohonsky, Terenzi et al. 2015) were maintained at 27°C, under 68% relative humidity and a 12/12h light/dark cycle. Mosquito larvae were reared in deionized water supplemented with minerals and fed on TetraMin Baby-E fish food from the day of hatching to the fourth larval instar supplemented with pieces of cat food. Male and female adults were provided free access to a 10% wt/vol sucrose solution for the first five days post-emergence (PE). Female mosquitoes were fed for 30 min on the blood of anesthetized mice. Adult progeny from Gal4-UAS crosses were maintained at 28°C.

### Plasmid construction

In a first step, the 3’ terminus of EYFP was removed from the plasmid pGEM-T[3xP3-eYFPafm-attB] (Adolfi, Pondeville et al. 2018) by *BsrG1-XbaI* digestion and replaced by a 3’ terminus of eYFPnls amplified by PCR from pSL[UAS-eYFPnls-g] (Lynd and Lycett 2012) and digested by *BsrG1* and *XbaI* to create the recipient plasmid pGEM-T[3xP3-eYFPnls-afm-attB]. The 1160 bp *hml* regulatory region from *D. melanogaster* was amplified by PCR from the *D. melanogaster* BAC genomic clone BACR02M20 (https://bacpacresources.org/). The reverse primer was designed to mutate an ATG located 79 bp before the *hml* start codon to AAG and therefore suppressing a putative upstream ORF. The *NotI-hml-EcoRI* PCR fragment was further cloned into psL[LRIM-gal4] (Lynd and Lycett 2011) after removal of the LRIM promoter with the same enzymes to give psL[*hml*-gal4]. The *hml*-gal4-SV40 cassette was then digested using *AscI* and *FseI* to be sub-cloned into the recipient plasmid pGEM-T[3xP3-eYFPnls-afm-attB]. The *hml* promoter was then amplified from psL[*hml*-gal4] with *FseI* forward and *PmeI* reverse primers. The gal80ts-SV40 cassette was amplified from pCASPER[*DEST-gal80ts*] (gift of Jacques Montagne) with *PmeI* forward and *FseI* reverse primers. Both PCR fragments were digested by *FseI* and *PmeI* and further cloned into pGEM-T[3xP3-eYFPnls-*hml*-gal4-attB] to create pGEM-T[3xP3-eYFPnls-*hml*-gal4-*hml*-gal80ts-attB]. This plasmid was sequenced to verify sequence integrity.

### hml-gal4 line establishment

The transgenic line was created by injecting 2387 embryos of the E phiC31 docking line (Meredith, Basu et al. 2011) with 250 ng/μl of *hml*-gal4 plasmid and 800 ng/μl of mRNA encoding an insect codon optimized mutant phiC31 integrase (Franz, Jasinskiene et al. 2011). Out of the 510 G0 larvae that survived (21.4%), 175 showing transient fluorescence (34.3%) were kept for further rearing. Recovered G0 adults (72 females and 71 males) were backcrossed to mosquitoes of the E line (pools of 10 G0 males and 50 females; all G0 females with 5x males) and all females blood fed. Among the progeny of these backcrosses, five transgenic G1 larvae were obtained (one from the G0 female pool of and four from the G0 male pools) and used to establish a homozygous transgenic *hml*-Gal4 driver line. Considering half of G0 individuals were sterile, the transformation efficiency is estimated ranging from 2.8% (if only one out of the four G0 males produced transgenic gametes) to 7% (if the four G0 males produced transgenic gametes). Preparation of the injection mix, microinjections, screening and stable homozygous line generation were carried out as previously described (Pondeville, Puchot et al. 2014). Correct site-specific integration was confirmed by PCR across the resulting *attL* and *attR* using specific primers (Meredith, Basu et al. 2011). Backcross to docking strain confirmed normal Mendelian inheritance of the eYFPnls marker.

### Bacteria and beads injection

To obtain bacteria expressing GFP, *E. coli* bacteria (Sure 2, Agilent Technologies) were transformed with pFPV25.1 (Valdivia and Falkow 1996) (Addgene plasmid # 20668) and grown overnight in a shaking incubator at 37°C in Luria-Bertani’s rich nutrient medium (LB broth) supplemented with ampicillin. After centrifugation, bacteria were resuspended in LB broth and cultures were normalized to OD_600_ = 0.5 using a spectrophotometer prior to injection. Yellow-green fluorescent FluoSpheres beads (1 µm diameter, Molecular Probes) were mixed with PBS to a final concentration of 0.08% solids per volume prior to injection. Cold-anesthetized 5-6 day-old females were injected into their thorax using a nanoinjector (Nanoject II, Drummond Scientific) with 69 nL of bacteria or 138 nL of fluorescent beads. Injected females were maintained with 10% (wt/vol) sucrose at 27°C until hemocyte perfusions three days after bacteria injection or one day after beads injection.

### Infection of mosquitoes with ONNV

The ONN-eGFP virus stocks were produced as described previously (Carissimo, Pondeville et al. 2015) from an infectious clone tagged with GFP in a duplicated subgenomic promoter (Keene, Foy et al. 2004). Female mosquitoes were allowed to feed for 15 min through a Hemotek membrane (Hemotek) covering a glass feeder containing the blood/virus mixture maintained at 37 °C. The infectious blood meal was composed of a virus suspension diluted (1:3) in washed rabbit blood and resuspended at 50% (vol/vol) in dialyzed rabbit serum (R4505; Sigma). ATP was added to a final concentration of 5 μM. The final blood-meal titer fed to mosquitoes was between 1–3 × 10^7^ pfu/mL. Non-infected females were treated the same way except that media used to produce the virus in cell culture was used instead of the virus suspension. Fully engorged females were transferred to small cardboard containers and maintained with 10% (wt/vol) sucrose at 28 ± 1 °C until hemocyte perfusions.

### Hemolymph perfusions

To collect circulating hemocytes, the last segment of the abdomen was cut, and mosquitoes were then injected into the thorax with phosphate buffered saline (PBS, pH 7.0) using a glass capillary mounted on a syringe. For RT-PCR experiments, PBS diluted hemocytes were collected in tubes on ice and centrifuged at 2000 rpm for 15 minutes at 4°C. Supernatant was gently removed before adding TRIzol (Molecular Research Center). For immunostaining, hemocytes were perfused on slides (ibidi; 10 females per well). Slides were left in the dark for 10 or 45 minutes at 28°C before removal of PBS-T and fixation.

### RNA extraction and RT-PCR

RNA from perfused hemocytes in TRIzol reagent was extracted according to the manufacturer’s instructions, except that 1-Bromo-3-chloropropane was used instead of chloroform and DNAse (TURBO DNase, Ambion) treatment was carried out. Reverse-transcription (RT) was performed using the MMLV retro-transcriptase (Promega) from 500 ng of total RNA in a final volume of 60 µl. cDNA was aliquoted and further stored at −20°C until PCR using the DreamTaq Green DNA polymerase (ThermoFisher) according to manufacturer’s instructions (40 cycles). Sequence of specific primers for *S7, gal4, gal80ts* and *mCD8::cherry* are given in Table S1.

### Immunostainings of tissues and hemocytes

Abdomen were dissected and carefully cut along the cuticle. Perfused hemocytes and abdomen tissues were fixed at room temperature (RT) for 20 minutes in 4% (w/vol) paraformaldehyde (PFA) diluted in PBS. Hemocytes were washed (three times, 15 minutes each at 4 °C) in PBS before overnight staining with DAPI 1X (405 nm, Sigma) and Phalloidin 1X (488 nm, Sigma or 647 nm, Cell Signaling) diluted in PBS at 4°C. After PBS washes (three times, 15 minutes each at 4 °C), mounting medium (ibidi) was used to replace PBS in wells. Tissues were treated the same way but PBS-Tween 0.05% was used instead of PBS. Tissues were mounted between slide and coverslip (24 mm × 24 mm) with an imaging spacer (1 well, diameter × thickness: 13 mm × 0.12 mm Grace Bio-Labs SecureSeal imaging spacer, Sigma-Aldrich) using mounting medium (ibidi). For mCD8 and PPO antibodies staining, fixed and washed hemocytes were blocked for at least 30 minutes in blocking solution (PBS-T 0.05%, 5% FCS [vol/vol], 5% BSA [w/vol], 0.05% Triton X-100 [vol/vol]) at 4°C. Hemocytes were incubated at 4°C overnight with a rat anti-mCD8 antibody (Ancell) diluted 1:100 or a rabbit anti-PPO2 (Fraiture, Baxter et al. 2009) at 1:1000, or a rabbit anti-TEP1 (Levashina, Moita et al. 2001) at 1:300 in blocking solution. Samples were washed (five times, 15 minutes each at 4°C) in PBS-T 0.05% and incubated with either an Alexa Fluor 488 goat anti-rat IgG (Thermo Fisher Scientific) or an Alexa Fluor 488 goat anti-rabbit IgG (Thermo Fisher Scientific) diluted 1:1000, DAPI 1X (405 nm, Sigma) and Phalloidin 1X (647 nm, Cell Signaling) in blocking solution for 2 h at RT. Three washes in PBS were carried out. Mounting medium (ibidi) was used to replace PBS in hemocyte wells. All steps were carried in the dark to preserve the endogenous mCD8::Cherry fluorescence. Images were acquired on an inverted Zeiss Observer Z1 (Axio vision software) for hemocytes, on a Leica (SP8) confocal microscope for tissues and processed with ImageJ and Adobe Photoshop.

## Ethical compliance

This study complied with all relevant ethical guidelines and regulations. Project (n° 2013-0132) approved by the Ministère de l’Enseignement Supérieur et de la Recherche – Direction Générale pour la Recherche et l’Innovation – Secrétariat « Autorisation de projet » – 1, rue Descartes, 75231 PARIS cedex 5. Human blood was obtained from ICAReB Platform, Center for Translational Research, Institut Pasteur, Paris, France.

## Data availability

All data are available upon request.

## Acknowledgments

We thank Frank Schnorrer for supply of pUAS-mCD8::Cherry plasmid, Amy Lind and Gareth Lycett for pSL[UAS-eYFPnls-g] and pSL[UAS-eYFPnls-g] and useful discussions, Ernst Wimmer for pBac[3xP3] and pslfa1180, Michele Calos pour pattB, Francois-Xavier Camval for pFPV 25.1 and Anthony James for phiC31 integrase optimized mutant plasmid template. We are also grateful to Jacques Montagne for confocal access.

## Supplementary Material

### *Hemolectin* orthology assessments across arthropods

Identification of putative orthologs of the *Drosophila melanogaster hemolectin* gene in arthropod species with sequenced and annotated genomes proceeded by searching genomic resources hosted at VectorBase release VB-2019-08 (Giraldo-Calderón et al.) and the OrthoDB hierarchical catalog of orthologs versions 9 and 10 (Zdobnov et al.). First, VectorBase was searched using the FlyBase gene identifier for *Hemolectin* (FBgn0029167), to identify the GeneTree (family of homologous genes) of which it is a member, GeneTree VBGT00190000010751. GeneTrees are built automatically by VectorBase using the EnsemblCompara pipeline (Vilella et al.) that uses the longest protein-coding translation of each gene with searches of the TreeFam (Schreiber et al.) library of sequence profiles followed by multiple sequence alignments with combinations of various aligners and then tree building with TreeBeST (Vilella et al.). This revealed orthologs in other Brachycera flies (*Musca domestica* and five *Glossina* species), three non-holometabolous insects (*Rhodnius prolixus, Cimex lectularius*, and *Pediculus humanus*), and the tick, *Ixodes scapularis* (Figure S3), as well as potential orthologs in two sandflies (incomplete gene models make orthology difficult to confirm with confidence). Second, the *D. melanogaster* Hemolectin protein sequence was searched using tBLASTn against all nucleotide data (genomes, transcriptomes, ESTs, etc.) from all species hosted by VectorBase. These searches confirmed the results from the automated GeneTree ortholog identification, i.e. several matches to mosquito genes that encode von Willebrand factor domains but no *Hemolectin* orthologs in mosquitoes. For a broader perspective across arthropods, searches of OrthoDB identified *Hemolectin* orthologs in many different groups including additional non-mosquito Diptera, as well as in Lepidoptera, Coleoptera, Hymenoptera, Hemiptera, Thysanoptera, Isoptera, Blattodea, Ephemeroptera, Odonata, Diplura, Collembola, Crustacea, Myriapoda, and Arachnida, but no orthologs in any mosquito species. These analyses strongly support the evolutionarily-rare loss of the *Hemolectin* gene from the genome of the last common ancestor of mosquitoes.

**Figure S1.**
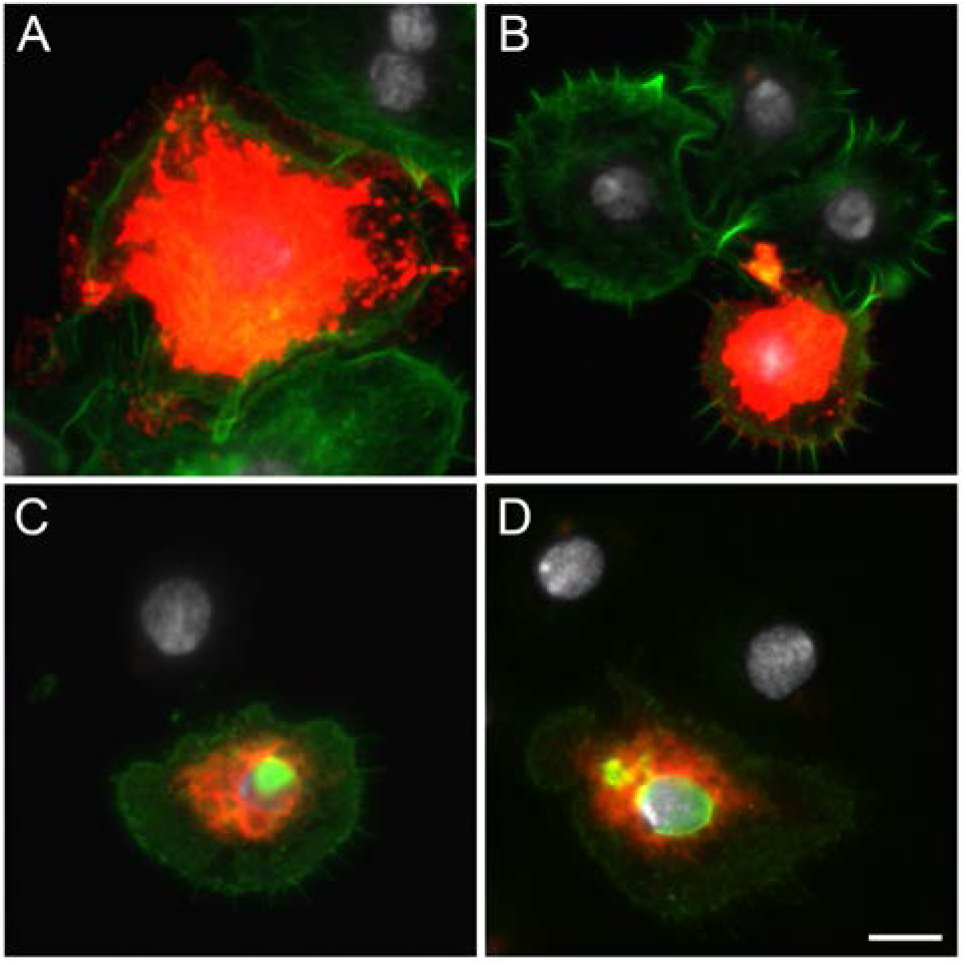
Hemocytes collected from *hml*>mCD8::Cherry blood fed females. (A, B) Over exposition of pictures I and J (Figure 2) showing Cherry at the cell membrane. Red: endogenous mCD8::Cherry; Green: Phalloidin; White: DAPI staining. (C, D) Anti-mCD8 antibody staining. Red: endogenous mCD8::Cherry; Green: anti-mCD8; White: DAPI staining. Scale bar is 5 µm.

**Figure S2.**
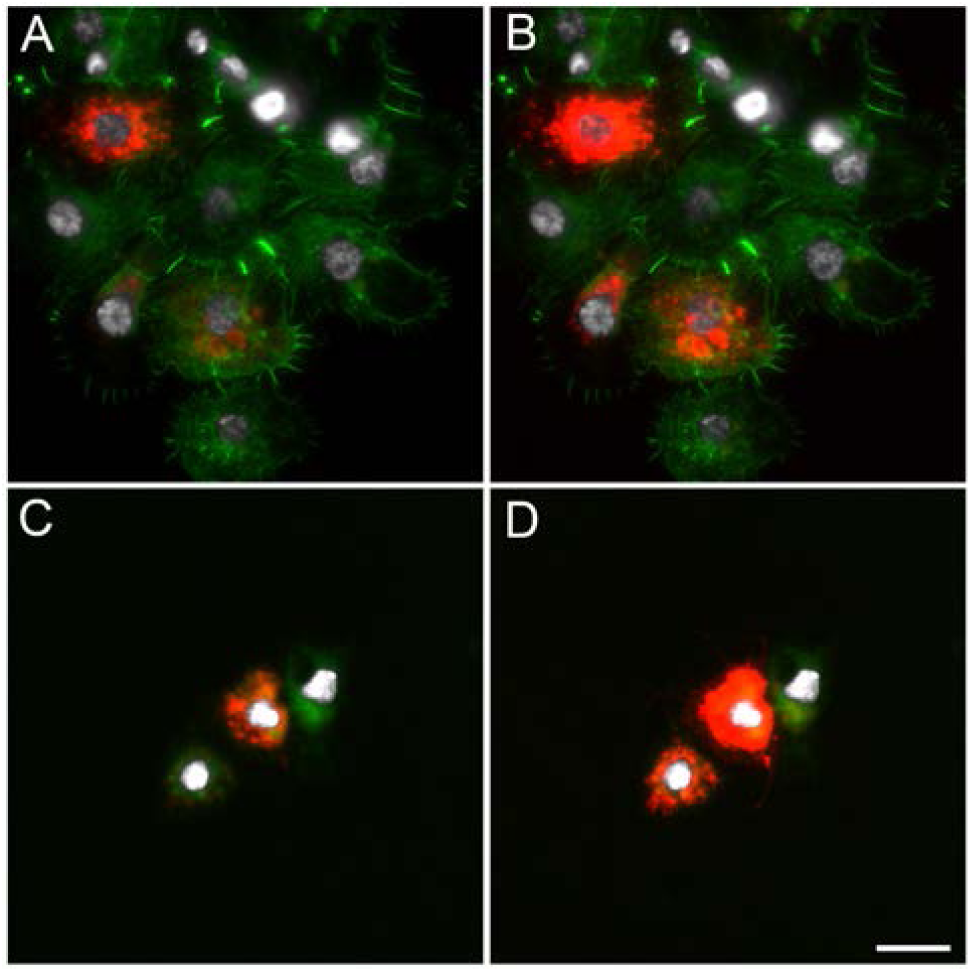
Hemocytes collected from *hml*>mCD8::Cherry blood fed females. B and D are highly exposed pictures from A and B respectively. Red: endogenous mCD8::Cherry; Green: Phalloidin; White: DAPI staining. Scale bar is 10 µm.

**Figure S3.**
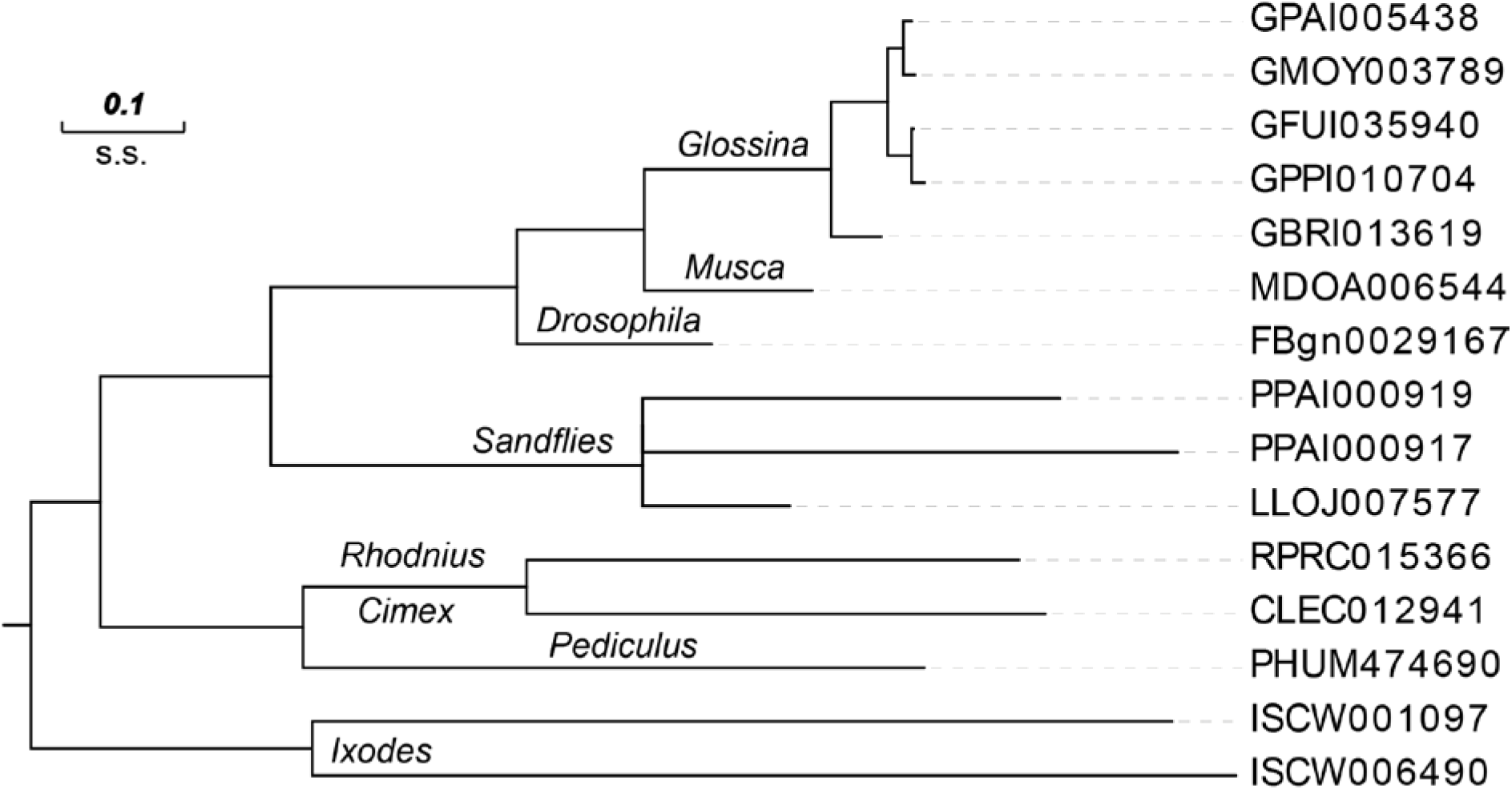
*Hemolectin* gene tree from VectorBase. Orthologs of *Drosophila melanogaster hemolectin* were found in other Brachycera flies and outgroup non-holometabolous insects and the *Ixodes* tick. Note that the two *Phlebotomus papatasi* genes are likely fragments of a single gene, similarly for the two *Ixodes scapularis* genes. No orthologs were found in any of the mosquito species at VectorBase.

**Table S1.**
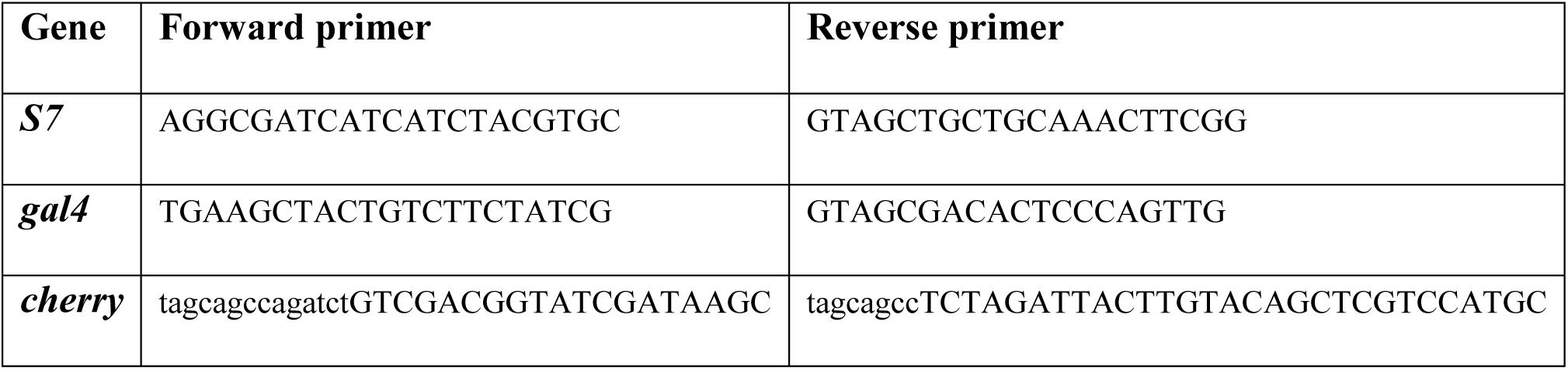
Sequence of primers used in the study.

